# Single-Cell and WGCNA Integrative Analysis Reveal the Key Chondrocytes Niches and Pathogenic Genes in Intervertebral Disc Degeneration

**DOI:** 10.1101/2025.03.07.642051

**Authors:** Yuyang Zhang, Xiaohui Wang, Dingjun Hao

## Abstract

Intervertebral disc degeneration (IDD) is a leading cause of low back pain, necessitating a comprehensive understanding of the molecular mechanisms driving disc degeneration to develop effective preventive and therapeutic strategies. Chondrocytes, as the predominant cellular constituents of the nucleus pulposus, are thought to play a pivotal role in the pathogenesis of IDD. In this study, we leveraged publicly available single-cell RNA sequencing (scRNA) data from the Gene Expression Omnibus (GEO) database for a comprehensive analysis. Employing unsupervised clustering methods, we successfully identified five distinct subtypes of chondrocytes and characterized two distinct terminal cell fates within the chondrocyte population. Through weighted gene co-expression network analysis (WGCNA), we identified RPL17, RPL13A, RPS18, and RPS24 as hub genes significantly associated with disc degeneration compared to control discs. Furthermore, our analysis of cell communication unveiled the activation of pro-inflammatory pathways in intervertebral disc degeneration. These findings shed light on the intricate complexity of chondrocyte involvement in the development of intervertebral disc degeneration and enhance our understanding of the role of chondrocyte niches within the nucleus pulposus during degenerative processes. Importantly, our study paves the way for potential targeted therapeutic approaches to address IDD.

## 1 Introduction

Low back pain (LBP) is a common global healthcare challenge, which is highly associated with intervertebral disc degeneration (IDD). Although disc degeneration is often asymptomatic, it sometimes causes sciatica and disc prolapse or herniation. IDD changes the height of the discs and the mechanics of the remainder of the spinal column, potentially altering the behavior of other spinal components such as muscles and ligaments. Besides, it can develop into spinal stenosis, a significant source of back pain and impairment in the elderly; its prevalence is increasing exponentially with the increasing age of people.

Intervertebral disc, a heterogeneous tissue, is made up of a gel-like nucleus pulposus (NP), a highly fibrous annulus fibrosus (AF), and cartilage endplate (CEP). The nucleus pulposus consists of an amorphous, water-rich extracellular matrix with sparse chondrocyte-like nucleus pulposus cells. The degeneration of NP is characterized by increased expression of matrix metalloproteinases and inflammatory agents (1). In addition, increased cell replication could cause cell cluster formation and cell death, which can ultimately result in cell senescence (2). The process eventually affects the AF and causes microtrauma and discomfort. Dysfunction of cells located in the intervertebral disc is responsible for all these changes. The intervertebral disc as an organ has a very limited capacity for intrinsic regeneration, likely because of an issue with the chondrocyte’s homeostasis maintenance. Investigating the chondrocyte niches in degenerative NP offers a potent method for delaying and even preventing IDD development and has great potential for clinical use.

Single-cell RNA sequencing (scRNA-seq) is a technique that provides previously unattainable insight into cell-level transcriptomes. The scRNA-seq technology improves understanding of disease pathogenesis mechanisms and cell-to-cell interactions. Weighted correlation network analysis (WGCNA) is well-acknowledged to find key clusters of highly associated genes related to module eigengene, furthermore identify the key candidate biomarkers or potential therapeutic targets. Combining these two powerful tools make it possible to broaden our insight into chondrocyte niches and pathogenic genes in IDD.

In human NP, scRNA-seq has assisted in the identification of novel cell types, the discovery of putative processes, and the investigation of cell communication from various perspectives. Based on single-cell transcriptome data, we described an overview of chondrocyte niches in human NP. Comparing degenerative NP and healthy NP allows us to investigate the key driver and trajectory of chondrocytes to degeneration. Based on ligand-receptor interactions, this study of cell communication between individual cells in IDD may assist in the development of novel pathways involved in IDD.

## 2 Materials and Methods

### Data Acquisition and Processing

Single-cell sequencing data was obtained from the GEO database (GSE160756, GSE165722). To get a more convincing result, only degeneration grades over III were included for analysis. We used SCtransform for normalization and variance stabilization of single-cell RNA-seq data and then integrated the data with Seurat v4.1.1(3). The downstream dimension reduction and clustering was completed with Seurat. Cells with more than 10% mitochondrial transcriptome were filtered. FindClusters was employed for clustering and resolution parameter was set at 0.5. We then identified 5 chondrocyte clusters, which were highly expressed SOX9, MIA, ACAN, and COMP. These chondrocyte clusters were selected for further analysis. Differentially expressed genes (DEGs) between chondrocyte clusters were calculated within Seurat 4.11. Gene Ontology (GO) analysis for these DEGs was completed by online platform g: Profiler and Benjamini-Hochberg FDR was set as 0.05.

### WGCNA analysis

We performed cluster analysis on samples and the height threshold was set as 500. In order to ensure that the correlation between connection and power is larger than 0.9, the optimal power (soft threshold) value was set as 3. We created a scale-free network and a topology overlap matrix (TOM) using this soft threshold. Additionally, using the function hclust from hierarchical clustering for module discovery, we created a gene (tree graph) hierarchical clustering tree. Setting the minimum module size as 40 and mergeCutheight as 0.3 prevent creating an excessive number of modules. A heat map with 26 modules was created after analyzing the associations between the built-in modules and the attributes. Finally, we produced the scatter plot and conducted a correlation analysis based on the gene importance of the module membership with the degeneration group. The p-value was adjusted by Bonferroni. The turquoise and blue modules were found to have higher correlations with p adj <0.05. We further performed subsequent functional enrichment and analysis of these two modules.

### Functional Enrichment Analysis and Hub gene identification

We analyzed Gene Ontology (GO) and Kyoto encyclopedia of genes and genomes (KEGG) pathway enrichment using clusterProfiler (version 4.2.2)(4). In addition, we considered Pathways of Gene Set Overrepresentation as significant only with BH adjusted p-value smaller than 0.05. Hub Genes were identified by CytoHubba plugin from Cytoscape software(5).

### Pesudotime analysis

The Monocle3 package was employed to plot trajectories(6). Single-cell sequencing data expression matrix was used to generate a CellDataSet using Monocle3. After dimensionality reduction, the cells were partitioned into metagroups. To plot the trajectory and color the cells according to the subcluster type, a cell trajectory module was utilized. Aligning cells from different batches using Batchelor (7). CytoTrace helped to predict differentiation states and determine less differentiated chondrocyte clusters(8).

### Cell-cell interaction inference

CellChat performed cell-cell communication analysis build on ligand-receptor pairings database (9). The fraction of cells expressing the relevant genes and the mean gene expression were used to calculate the mean, which represents the ligand and receptor mean expression in a particular cell type. The proportion of the mean equal to or greater than the actual mean, describing the likelihood of a certain cell type for a particular receptor-ligand complex, determines the P-value. The results were visualized by the circlize (v0.4.15) R package(10).

### Transcription Factor Prediction

Transcription factor (TF) networks were built using scRNA expression matrices and the SCENIC (v1.2.4) R toolkit(11). Given the common expression of driving regulators and targets, the GENIE3 project (v1.16.0) constructed a regulatory network. AUCell and RcisTarget were used to prune target modules as well as assess regulatory network activity across all cells, respectively.

### Statistical Analysis

R software version 4.1.3 was used for all statistical analysis and display of findings. To compare two groups, the Wilcoxon test was utilized. BH and Bonferroni methods were applied to adjust p-value.

## 3 Results

### 3.1 Cell cluster identification of Human NP

In the purpose of comparing the healthy and degenerative NP at single-cell resolution, we employed Seurat v4 to integrate scRNA data from 2 separate GEO datasets. GSE160756 and GSE165722 is data from healthy adults and IDD patients, respectively, displayed in Table S1. To investigate the most representative patterns between healthy and degenerative NP, we excluded samples under degeneration grades III in GSE165722. SCT normalization method(12) was used for anchors detection and data integration and we obtained 11 clusters by a shared nearest neighbor (SNN) modularity optimization-based clustering algorithm. UMAP plot showed that the batch effect was removed after integration (Figure 1A). We used the “Seurat” package in R software to integrate data for downstream analysis. Uniform manifold approximation and projection (UMAP) was used to cluster and visualize all cells together based on their transcriptional similarity. Unsupervised UMAP analysis revealed 11 transcriptionally distinct subpopulations (Figure 1B).

**Figure 1.**
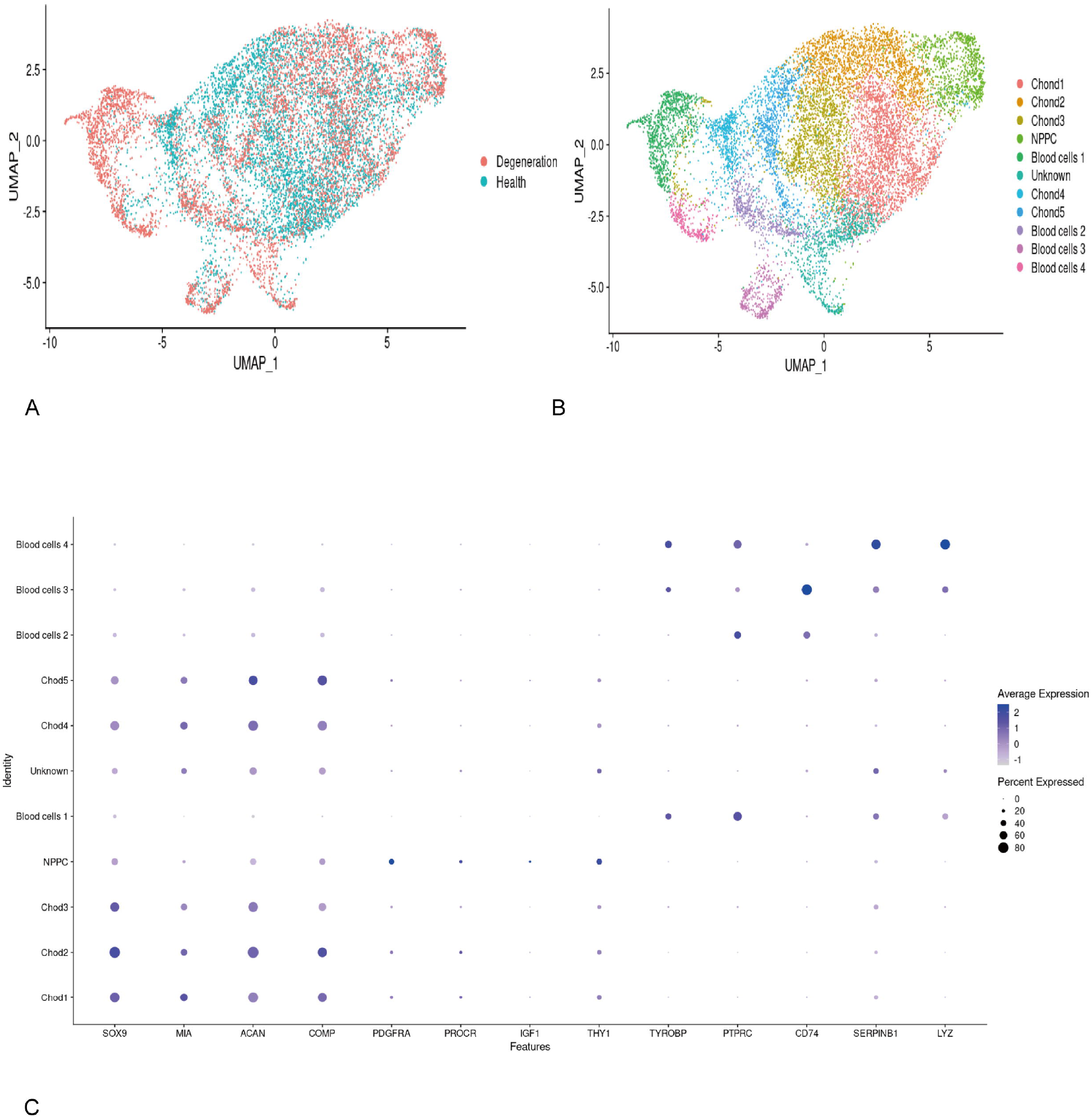
Cell cluster identification of Human NP. **(A)** UMAP showed the batch effects had been removed after integration. **(B)** After integration, cells from NP could be divided into 11 subsets. **(C)** Dotplot for identity markers from clusters 1-11

A total of 12000 cells with 20267 features were included for analysis after pre-processing data and quality control to investigate the most representative patterns between healthy and degenerative NP, we excluded samples under degeneration grades III in GSE165722. SCT normalization(12) was used for anchors detection and data integration and we obtained 11 clusters by a shared nearest neighbor (SNN) modularity optimization-based clustering algorithm.

Unsupervised clustering illustrated three major groups in nucleus pulposus cells based on specific gene patterns: NPPC (specifically expressing PDGFRA, PROCR, IGF, THY1), chondrocytes (specifically expressing SOX9, MIA, ACAN, and COMP(13-15)), and blood cells (specifically expressing PTPRC, CD74, SERPINB1, LYZ, and TYROBP) (Figure 1C). Five chondrocyte clusters were identified and selected for further analysis.

### 3.2 Biological Function of Chondrocyte clusters

Chondrocyte secretes extracellular substances, including proteoglycans and collagens required for homeostasis of NP. After identifying differentially expressed genes, we used Gene Ontology (GO) analysis to further understand the biological function and heterogeneity of chondrocyte subsets (Table S2). Of note, genes including CHI3L1, CHI3L2, and APOD promote Chond1’s multifunctionality of positive regulation of catabolic process, metabolic process, and lipoprotein oxidation (Figure 2A). Collagen fibril organization and extracellular matrix structural constituents were considerably enhanced in Chond2 (Figure 2B). Our results revealed that Chond3 clusters were characterized by regulations of transcription and cell death (Figure 2C) while Chond4 were classified as inflammatory response subsets and ECM function (Figure 2D). IGFBP1, COL1A1 and CFLAR were more active in Chond5, which were described for cellular response to biological process (Figure 2E).

**Figure 2.**
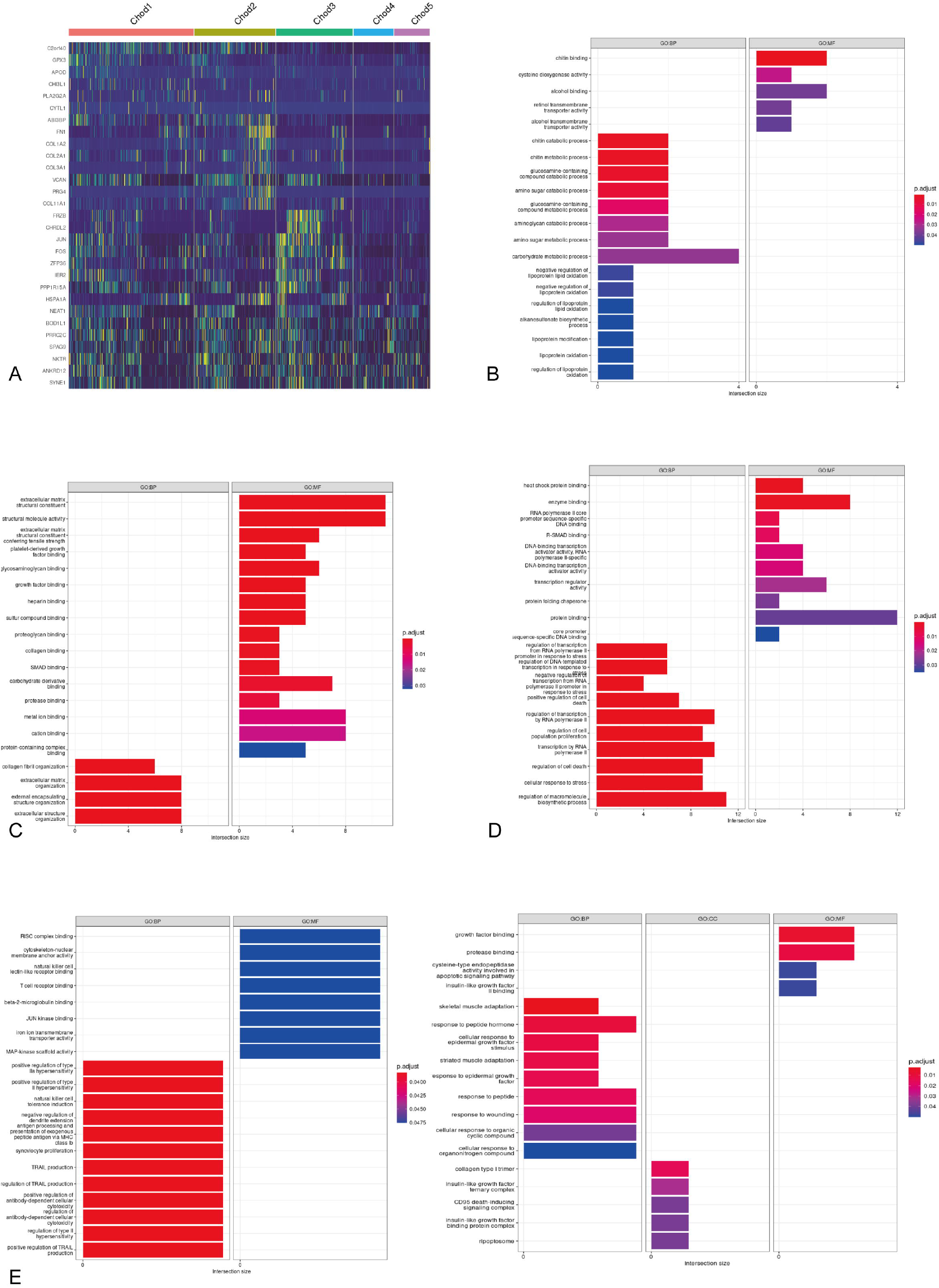
Biological Function of Chondrocyte clusters. **(A)** Chondrocyte cluster 1 was mainly associated with metabolic process. **(B)** GO enrichment annotation for chondrocyte cluster 2 revealed to be involved in ECM. **(C)** Chondrocyte cluster 3 related to regulations of transcription and cell death. **(D)** Chondrocyte cluster 4 were charactered by inflammatory response and ECM function. **(E)** GO enrichment annotation for chondrocyte cluster 5.

### 3.3 Identification of key modules by WGCNA

Weighted gene co-expression network analysis (WGCNA) could find clusters (modules) of highly correlated genes, summarize these clusters using module eigengene, and associate modules with external sample features (16). The indices of scale-free fit for calculated soft power threshold levels are shown in Figure 3A. We performed sample clustering analysis after the ideal soft power was set as 3.

**Figure 3.**
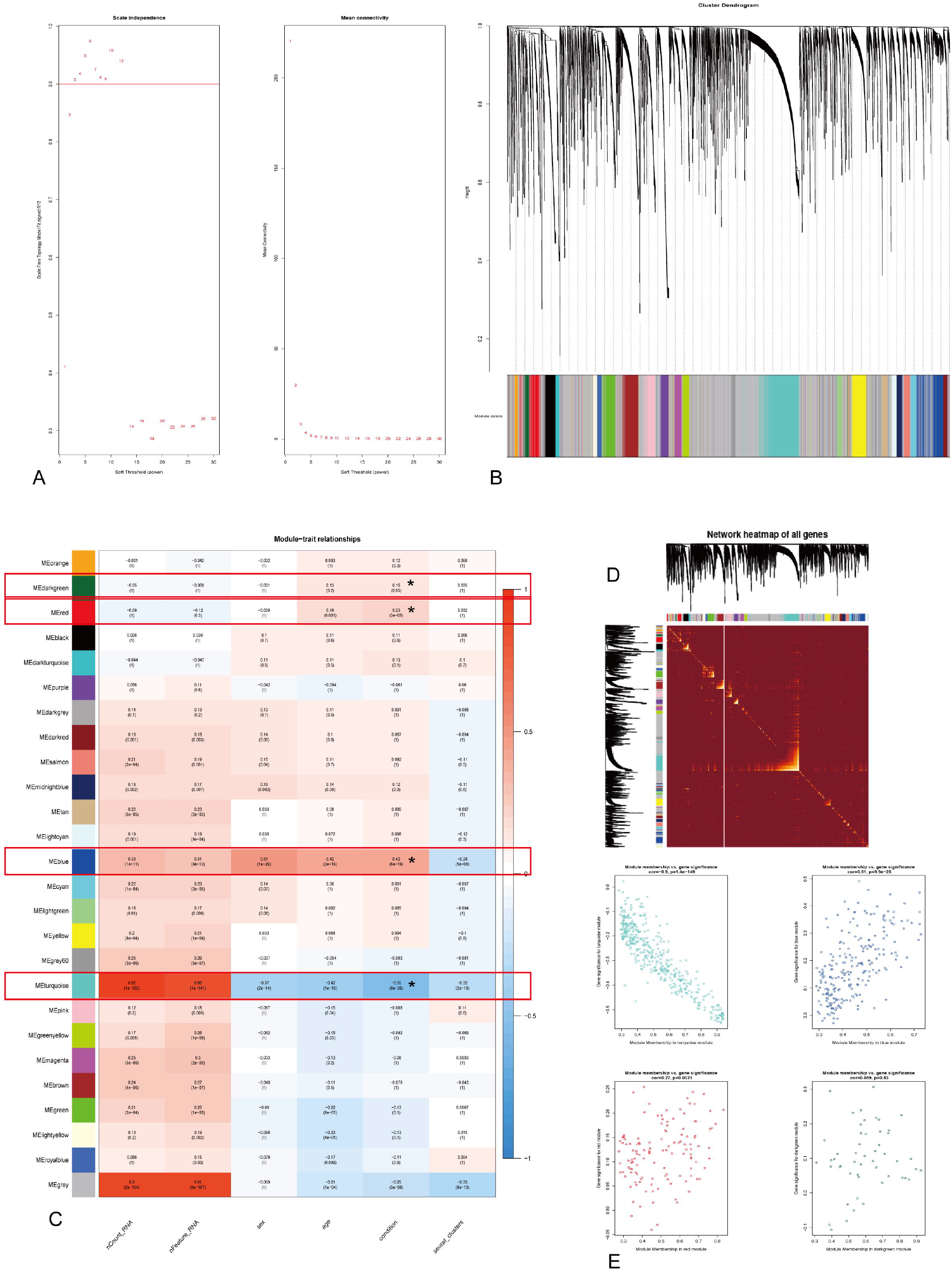
Identification of key modules by WGCNA. **(A)** Scale-free plots determined the best soft-power of WGCNA as 3. **(B)** Dendrogram of identified modules. **(C)** Correlation heatmap showed dark-green, turquoise, red, and blue modules strongly correlated to IDD traits. **(D)** Network TOM heatmap plot. **(E)** Among the four selected modules, blue and turquoise had the highest correlations.

We grouped the adjacency matrix into a TOM matrix and used hierarchical clustering to generate a module tree predicted on the clustering of hclust hierarchy. For module identification, a dynamic tree pruning method is applied. The cutting height of the module merging dendrogram is set to 0.3, and the minimum module size for module detection is set to 40 (Figure 3B). Correlation analysis is performed between the various modules and feature categories produced by the clustering. A topological overlap matrix (TOM) was used to hierarchically cluster metagenes from scRNA samples (Figure 3D).

Twenty-six modules, including grey modules which meant genes failed to be clustered, were identified. The heat map listed the p-values and correlation coefficients for each module and each characteristic (Figure 3C). Four key modules with significance of condition were identified. Among these four significant modules, turquoise and blue modules had larger correlation coefficient with condition factor (Cor =0.55, 0.42, p-value =9e-36, 6e-19 respectively). The correlation between module membership in modules and gene significance in the trait of degeneration was shown in Figure3E. The correlation in turquoise was calculated to be 0.9, p-value = 1.4e-149 and in blue module was revealed to be 0.61, p-value = 9.9e-25. Taken together, genes in the turquoise and blue modules had optimal positive associations with the degeneration. Genes in the turquoise and blue modules were then subjected to functional analysis.

### 3.4 Enrichment analysis of GO, KEGG, and Hub genes identification

The turquoise and blue module has 640 genes in total. We then performed enrichment analysis for these identified genes. There were 226 GO terms identified and 178 were biological processes, 30 were cellular components, and 18 were molecular functions among them. Most of them are involved in protein folding and binding, extracellular matrix structural constituents, ribosome, and cytoplasmic translation. The MAPK signaling pathway, ribosome, and oxidative phosphorylation were recognized in the KEGG pathway enrichment study (Figures 4A-D). By using cytoHubba plugin in Cytoscape software (MNC top 15), we identified the following top 15 ranked hub genes (Figure 4E). Most hub genes contribute to function of ribosomes.

**Figure 4.**
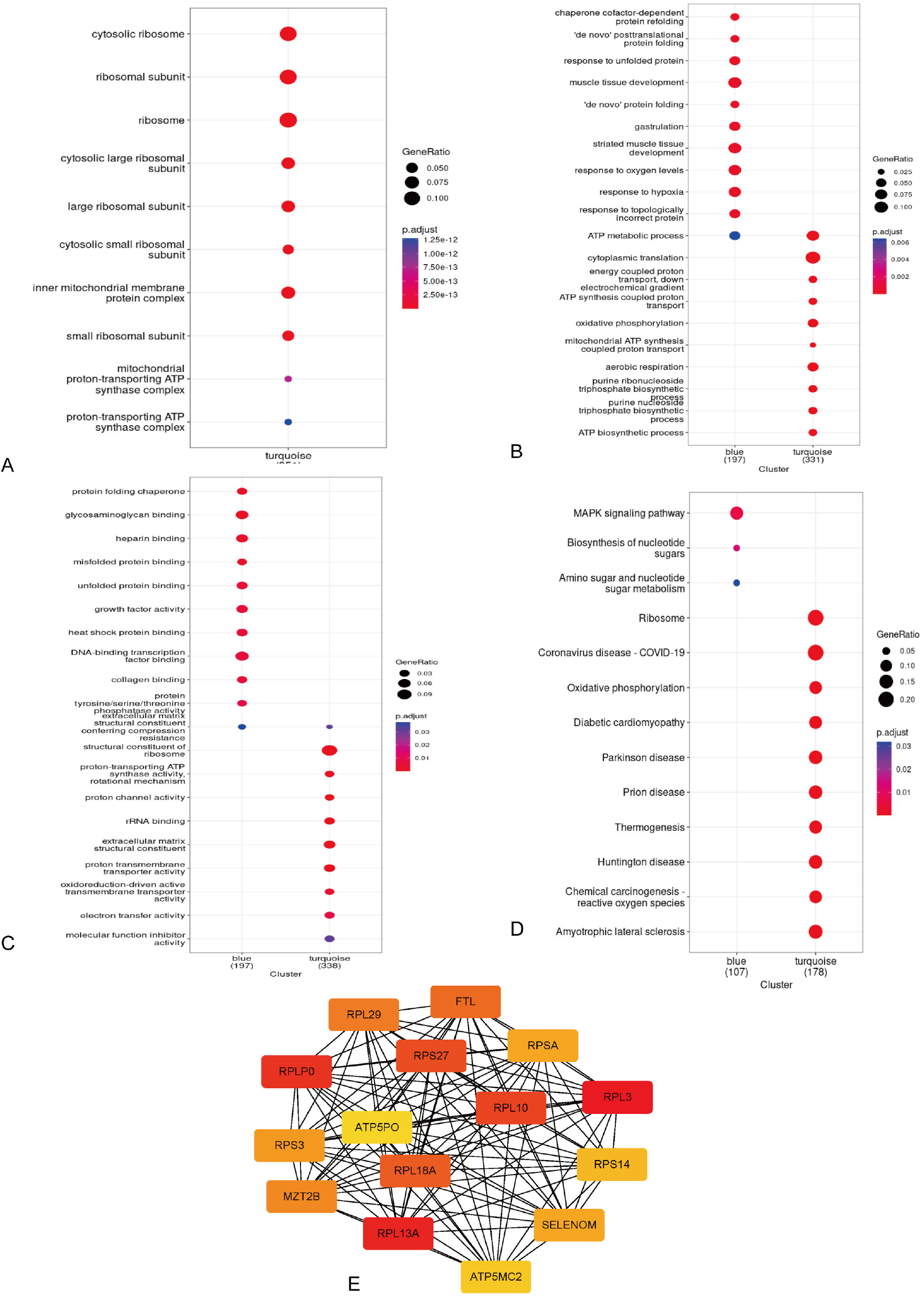
Enrichment analysis of GO, KEGG, and Hub genes identification CC BP MF. **(A-C)** The GO terms of the Cellular Component (CC), Biological Process (BP), and Molecular Function (MF) categories enrichment of blue and turquoise modules identified by WCGNA. **(D)** Annotated KEGG pathway of blue and turquoise modules. **(E)** Top 15 hub genes from blue and turquoise modules determined by CytoHubba plugin.

### 3.5 Transcriptional Regulation in Chondrocyte Subpopulations

Transcription of cells is determined by gene regulatory network (GRN), in which several transcription factors (TFs) and cofactors interact with each other and regulate their downstream target genes. To further understand the molecular mechanism of IDD, we employed SCENIC package to predict the transcriptional regulation in chondrocyte subpopulations. Regulons in SCENIC are referred to for modules with significant motif enrichment of the correct upstream regulator.

The regulon activities for different chondrocyte clusters were shown in the heatmap (Figure 5A). We then divided these cells by different conditions and found that healthy intervertebral disc had higher activity of CEBPD, ID1 JUN, JUNB, and FOS while MAFG, VDR, and EP300 and was more activated in degeneration state (Figure 5B).

**Figure 5.**
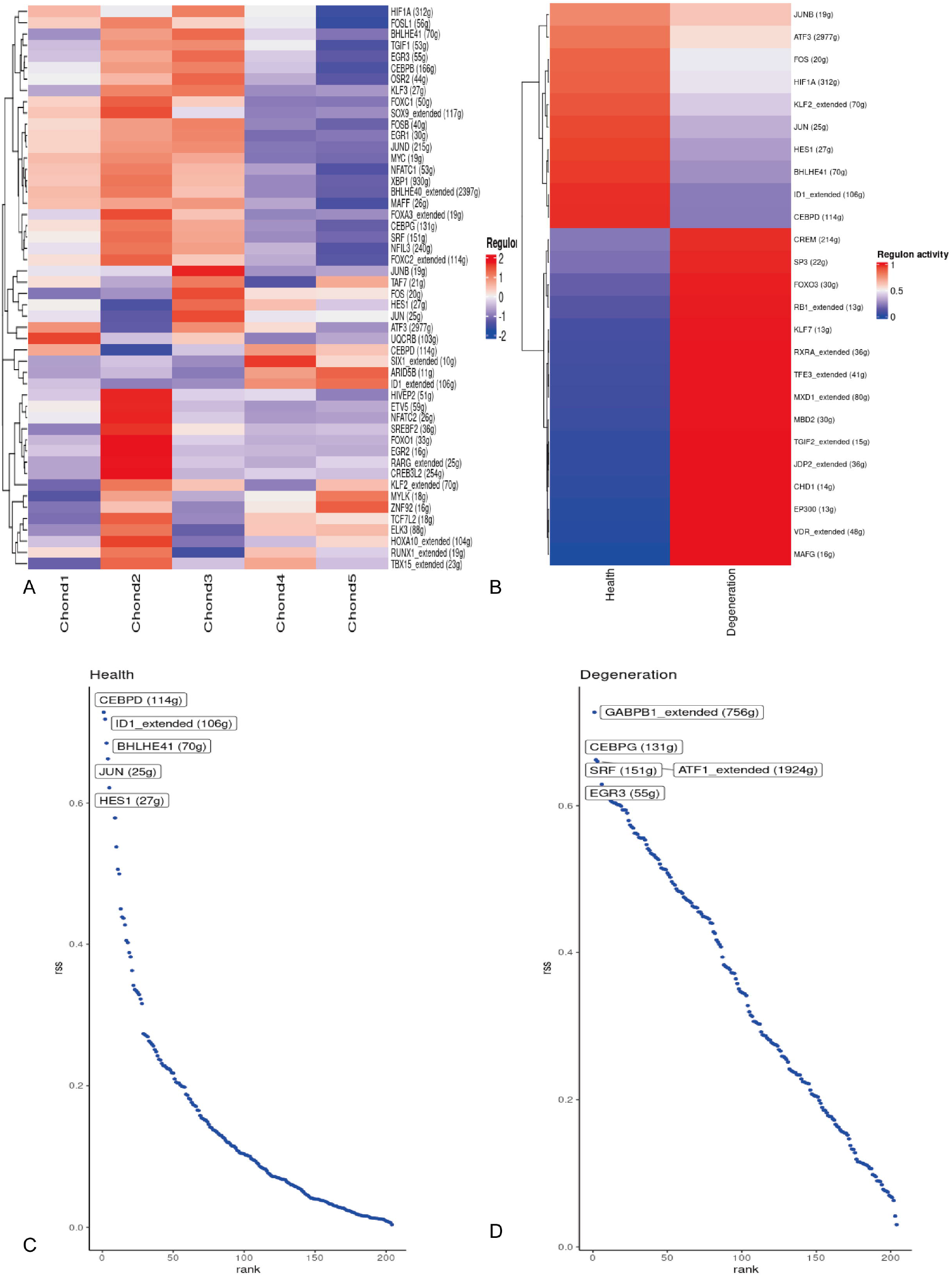
Transcriptional Regulation in Chondrocyte Subpopulations. **(A)** Heatmap showed activities of regulons from different chondrocytes. **(B)** Dendrogram of identified modules. **(C)** Correlation heatmap showed dark-green, turquoise, red, and blue modules strongly correlated to IDD traits. **(D)** Network TOM heatmap plot. **(E)** Among the four selected modules, blue and turquoise had the highest correlations.

We then employed Regulon Specificity Score (RSS) to identify the condition-specific regulons. The most specific regulons in healthy state were CEBPD, ID1, BHLHE41, JUN, and HES1 while in degenerative state were GABPB1, CEBPG, ATF1, SRF, EGR3 (Figures 5C, D).

### 3.6 The Trajectory of Chondrocytes

Cells change from one functional “state” to another throughout development in response to stimuli. Different genomes are expressed by cells at different stages, resulting in a dynamic repertoire of proteins and metabolites that perform their functions. Transcriptional recombination occurs when cells switch to another state, with some genes repressed and others newly activated. IDD development and changes in expression patterns in chondrocytes are highly associated.

We used Monocle3 trajectory analysis to investigate chondrocyte transformation in NPs during IDD development (Figure 6A). CytoTrace was also employed to determine developmental trajectories (Figure 6B). Cells at the root of pseudotime were coloured in darkblue. Clusters of Chond4 and Chond1 have more potential for differentiation. Patterns of several chondrocyte subtypes were included for comparison in the pseudotime trajectory. Chond1 and Chond4 are mostly found near the root. Chon3 and Chond2 are primarily arranged at the terminal of cell fate (Figure 6C). Mostly Chond5 cells were not found in the trajectory.

**Figure 6.**
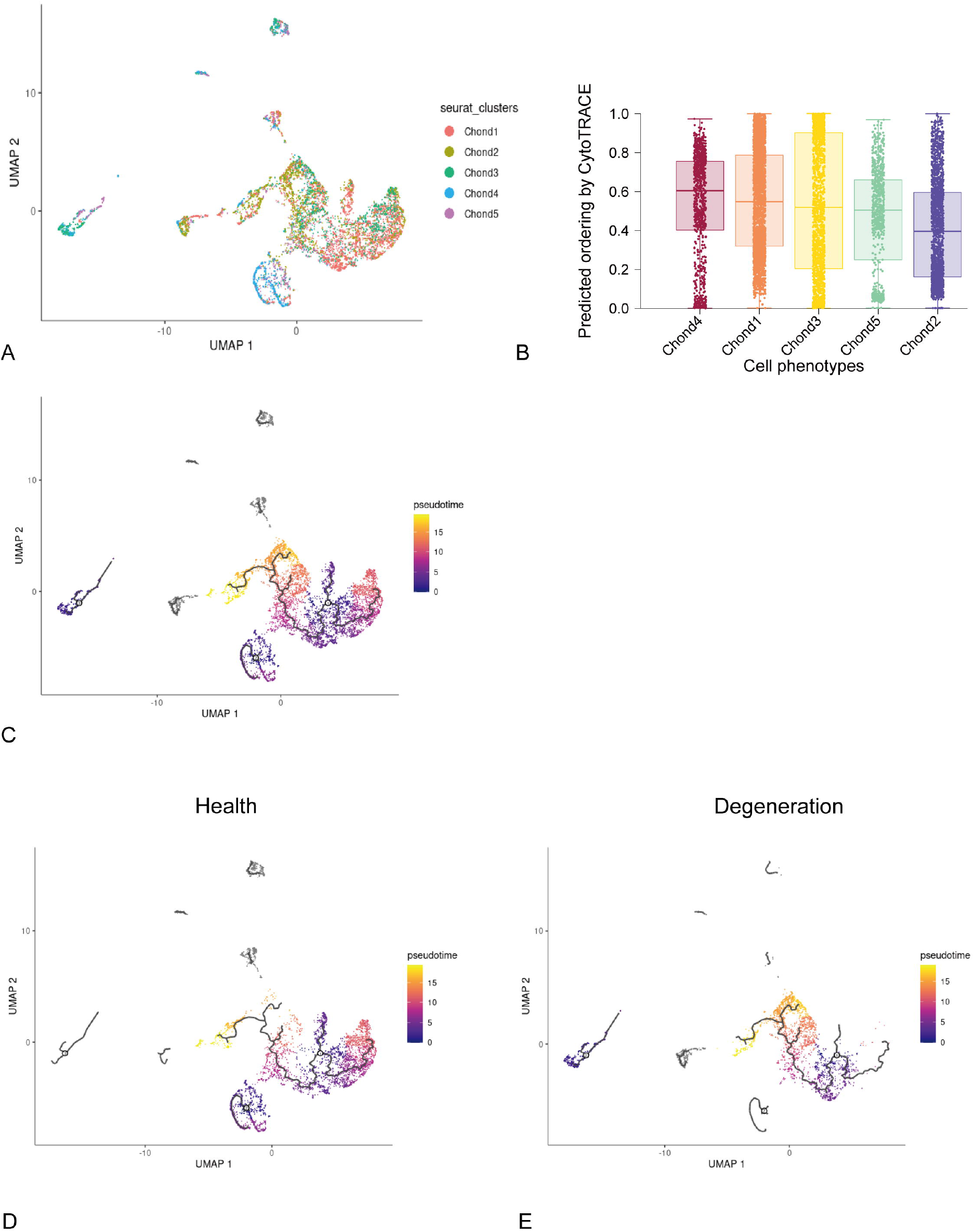
The trajectory of chondrocytes. **(A-C)** Pseudotime trajectories of all chondrocytes from integrated scRNA data. **(D, E)**, Comparison of cell fates of chondrocyte subsets in healthy and IDD state.

Further analysis revealed that the proportion of Chond2 and Chond3 in IDD progressively rose, as the disease progressed (Figures 6D, E). As shown in the enrichment analysis, Chond2 gene patterns were involved in collagen fibril organization and extracellular matrix structural constituent while Chond3 involved in cell death, as the disease progressed (17-19). The chondrocyte clusters distribution curve of differentiation fate was found to be compatible with the pathological process of IDD, according to our findings. Trajectory analysis indicated that chondrocytes differentiate into two cell fates during IDD. Investigation of chondrocyte differentiation may inspire effective approaches to treat IDD, as ECM alterations and cell death processes are significantly associated with both cell fates.

### 3.7 Constructing Cell Communication Network for NP

Ligand-receptor pairs promote or inhibit disease development, regulate cell function and fate, and allow cells to connect with each other. We performed receptor-ligand interaction-based analysis using CellChat to reveal cellular communication in NP. Non-degenerated NP cellular communication was defined as essential cellular communication that preserves NP function (Figure 7A). As shown in Figure 7B, cellular communication changed dramatically under degenerative state. Venne plot showed the predicted pathway between healthy and degenerative states. Of note, inflammation pathways including Annexin, IL1, and TNF pathways are specifically active in degenerative states (Figure 7C, Table S3), as described in previous literatures(20, 21). In addition to these pro-inflammatory pathways, IDD also exhibits highly active pathways such as antigen presentation pathway (MHC-I/ II), adhesion, and migration (Periostin, ICAM).

**Figure 7.**
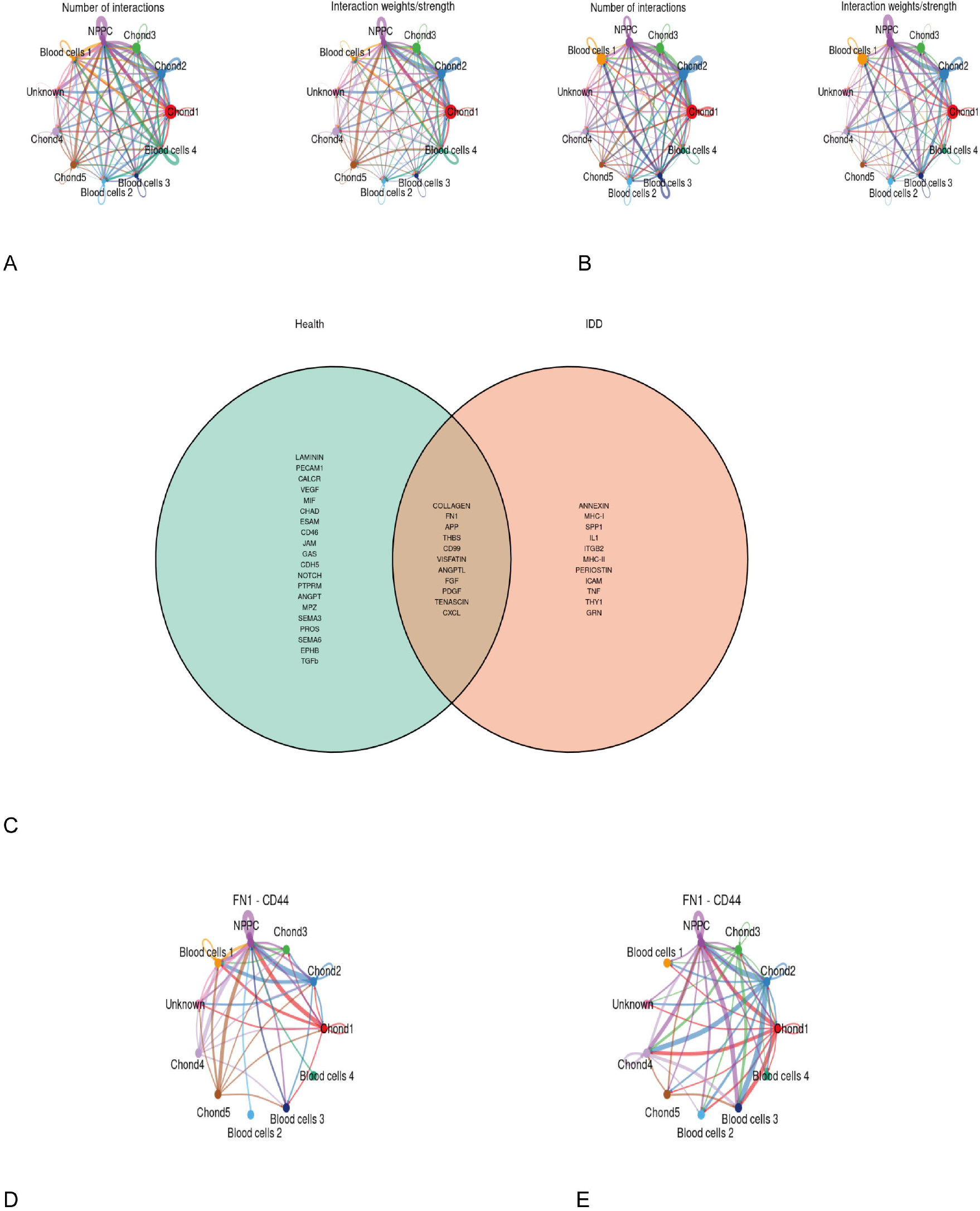
Constructing cell communication network. **(A)** Number of interactions and interaction strength in healthy NP. **(B)** Number of interactions and interaction strength in degenerative NP. **(C)** Venn diagram showed the pathways intersect under different condition. **(D)** FN1-CD44 pathway interactions in healthy NP. **(E)** FN1-CD44 pathway interactions in degenerative NP.

To investigate the cell communication disturbing during the development of IDD, we summarized 11 conserved types of cell communications separately (Figure 7C). The most noticeable change is that FN1-CD44 was highly active in degenerative NP. Chondrocyte fibronectin 1 (FN1) activates P13K/AKT to promote cartilage matrix production and chondrogenic differentiation (22). FN1 was reported significantly increased expression in degenerative disc disease(23), consistent with our results.

## 4 Discussion

Low back pain (LBP), which is a huge burden for health and social care systems, is prevalent in aging societies(24, 25). In the spine, the intervertebral discs are located between adjacent vertebrae. intervertebral discs allow the vertebrae to move slightly, act as ligaments that hold the vertebrae together, and act as shock absorbers for the vertebrae. Intervertebral disc degeneration (IDD) is characterized by the losing elasticity, flexibility, and absorption function in disc, as well as annulus fibrosus (AF) fragility(26). Low back pain due to disc degeneration is a major cause of population deterioration and loss of work capacity, with negative social and economic consequences (27). IDD is linked to higher levels of pro-inflammatory cytokines, such as TNF, IL-1α, IL-1β, IL-6, and IL-17(21). However, the current clinical-proven treatment strategies for IDD like analgesics, anti-inflammatory medications, and physical therapy ignore the prevention of IDD(28). Consequently, investigating mechanisms of disc degeneration is important for finding new prevention and treatment strategies.

It is generally accepted that cell components of nucleus pulposus are highly heterogeneous. From the histology slides, cells of the normal nucleus pulposus widely express Sox9 and collagen II mRNA that indicate a chondrocyte phenotype(29). This was also confirmed by high-resolution single-cell RNA sequencing that chondrocytes are majority of cells in nucleus pulposus (30). Of note, it was reported that collagen II expression, like aggrecan, decreased as the degree of degeneration increased(29). These previous results inspired us to think that chondrocytes in NP might play an important role in the development of IDD.

GO and KEGG enrichment analysis revealed that significant modules identified by WGCNA were associated with ribosome function, protein folding, oxidative phosphorylation, and MAPK signaling pathway. Previous studies have revealed that the MAPK/ERK signaling pathway regulates ECMis metabolism of IDD cells and oxidative stress are involved in IDD development (31). Oxidative stress was also reported to contribute to the progression of IVD degeneration(32). These evidences confirmed our results from WGCNA.

Our top 15 identified hub genes were dominated by ribosomal proteins coding genes, such as RPL3, RPL10, RPL13A, RPL18A, RPL29, RPLP0, RPS3, RPSA, and RPS14. Ribosome biogenesis and protein translation play an essential role in cell growth, differentiation, and proliferation. Few researches have been done on ribosomal proteins and their relationship to disc degeneration. In a bioinformatic study published in 2015, deregulation of ribosomal genes (RPL8, RPS16, and RPS23) was observed to be related to disc degeneration in comparison to scoliotic discs(33). Another bioinformatic study illustrated that RPL17, RPL13A, RPL18, and RPS24 were identified as the top 10 hub genes of the protein-protein interaction network implicated in degeneration compared to control discs(34). Liu Y et al established a mouse model deficient in binding RPL11 and RPL5, the transgenic mouse prone to fat accumulation under normal feeding conditions and hepatic steatosis under acute fasting conditions (35). One L13a allele disruption grants resistance to lipid-induced oxidative apoptosis(36). These studies indicate that ribosomal proteins might be involved in disc homeostasis as well as lipid metabolism.

Transcription factors regulate the activity and velocity of translation in cells. Interestingly, we found that the activity of several myogenesis and adipogenesis transcription factors (EGR3, ATF1, CEBP/γ) (37-39) was enhanced in degenerative chondrocytes. Recently, several studies have proposed the causative relationship between lipid dysfunction and degenerative disc development(40). A retrospective study of 790 Chinese patients illustrated that serum lipid levels were highly positively correlated with lumbar disc degeneration(41). In vitro study, Li et al observed that ox-LDL treatment can reduce the viability of nucleus pulposus cells(42).

The breakdown or storage of fats for energy and the creation of structural and functional lipids are both components of lipid metabolism, which involves the synthesis and degradation of lipids in cells. Although overweight and obesity are risk factors related to disc degeneration, the molecular mechanisms of the association remain speculative. It has been thought that overweight and obesity cause disc degeneration by getting excessive loadings. However, H Jonsson et al have reported elderly females with atherosclerosis have more prone to hand osteoarthritis than those without atherosclerosis(43). Such results refute the hypothesis that being overweight or obese only indirectly causes osteoarthritis through changed biomechanics due to greater loading. This behavior may be partly explained by secondary mediators generated by adipocytes, such as adipocytokines, which cause an inflammatory response, or proinflammatory cytokines and chemokines. Another possible mechanism is vascular insufficiency to the vertebrae caused by atherosclerosis or excessive serum lipids, which might reduce nutrition and metabolite transfer into the disc. Other proposed explanations include metabolic disorders or gene-environmental interactions.

Trajectory inference is a method to determine the pattern of a dynamic process in single-cell transcriptomics. Pseudotime is a measurement of how far an individual cell has progressed through a process cell differentiation. Usually, less differentiated cells are set as roots in the trajectory of pseudotime. Clusters of Chond4 and Chond1 were determined as roots of trajectory by an unsupervised method. Then chondrocytes in a degenerative state exhibit more tendency to differentiate into Chond2 and Chond3, which were characterized by ECM constituent and cell death, respectively. Besides, inferred cell communication illustrated that pro-inflammatory pathways are specially activated under a degenerative state, indicating the occurrence of inflammation during the development of IDD.

Finally, although this study identified the underlying mechanism of IDD and connected them with clinical traits, we must acknowledge our limitations. Available data lack sufficient general clinical data on patients, and only a few series contain weight information, making it difficult to incorporate various factors into the analysis

## 5 Conclusions

By using cell communication and trajectory analysis, we characterized pro-inflammatory and metabolic enhancement during the development of IDD, which might be driven by myogenesis and adipogenesis transcription factors. WGCNA illustrated ribosomal proteins might be involved in disc degeneration. This study broadens our understanding of IDD and reveals novel procedures, biomarkers, and potential therapeutic targets for the benefit of patients.

## 7 Author Contributions

Conceived the idea: DH and YZ; Manuscript draft: YZ; Downloaded and collected data: XC; Analyzed the data: YZ, XC, KS; Prepared figures: YZ, XW, ZC, and TY; Redressed the manuscript: XC, KS; Reviewed the manuscript: All authors. This study was completed with teamwork. Each author had made corresponding contribution to the study. All authors contributed to the article and approved the submitted version.

## 9 Acknowledgments

We unfeignedly thank Dr. Zhenggang Wang (Tongji Hospital, Huazhong University of Science and Technology) for generously sharing his experiences and codes.

## 1 Data Availability Statement

The datasets analyzed for this study can be found in the GEO database [https://www.ncbi.nlm.nih.gov/geo].

